# Pial collaterals develop through mosaic colonization of capillaries by arterial and microvascular endothelial cells

**DOI:** 10.1101/2023.10.09.561542

**Authors:** Tijana Perovic, Irene Hollfinger, Stefanie Mayer, Janet Lips, Monika Dopatka, Christoph Harms, Holger Gerhardt

## Abstract

Collaterals are unique blood vessels present in many healthy tissues that cross-connect distal-end arterioles of adjacent arterial trees, thus providing alternate routes of perfusion. Stroke patients with superior pial collateral flow respond better to treatments and present with an overall improved prognostic outcome. However, how pial collaterals develop in the embryo and how they reactivate upon stroke remains unclear. Here, using lineage tracing in combination with three-dimensional imaging, we demonstrate that mouse embryos employ a novel mechanism to build pial collaterals, distinct from their outward remodeling following stroke. Endothelial cells (ECs) of arterial and microvascular origin invade already existing pre-collateral vascular structures in a process which we termed mosaic colonization. Arterialization of these pre-collateral vascular segments happens concurrently with mosaic colonization. Despite having a smaller proliferative capacity, embryonic arterial cells represent the majority of cells that migrate to form nascent collaterals; embryonic microvascular cells, despite their higher proliferative potential, form only about a quarter of collateral endothelial cells. Moreover, postnatal collateral growth relies much more on self-replenishment of arterial cells than on microvascular contribution. Following ischemic injury, pial collateral outward remodeling relies on local cell proliferation rather than recruitment of non-arterial cells. Together, these findings establish distinct cellular mechanisms underlying pial collateral development and ischemic remodeling, raising the prospect for future research to identify novel, collateral-specific therapeutic strategies for ischemic stroke.

## Introduction

Ischemic stroke is one of the leading causes of death worldwide as it disrupts blood supply to the brain, retina, or spinal cord (Saini et al., 2021). While reperfusion can be achieved through thrombolytic or endovascular treatment, many patients do not respond well to these therapies (Ginsberg, 2018). Alternative therapies could focus on augmenting pial collateral blood flow, either by the induction of new collaterals or by functionalization of existing collaterals. Pial collaterals cross-connect neighboring cerebral arteries, creating a natural bypass capable of restoring blood flow to the ischemic region (Coyle and Heistad, 1991; Wei et al., 2001; Pipp et al., 2004; Okyere et al., 2020). Stroke patients with good collateral flow respond better to existing treatments, have a higher likelihood of major reperfusion, and display an overall better prognosis (Iwasawa et al., 2016). Although pial collaterals were anatomically described in the 17th century (Willis, 1664), functional imaging techniques have only recently clarified the importance of collateral grade on stroke outcome (Lin and Liebeskind, 2016). To date, many fundamental questions regarding pial collateral formation in development and their remodeling in the aftermath of ischemia remain unresolved.

Several mechanisms have been proposed to describe collateral formation in the mouse brain, heart, and hindlimbs. Of note, in the heart and hindlimb of mice, collateral formation is only induced by ischemia and not during development (Faber et al., 2014; Perovic et al., 2022). Collaterals present in the adult may arise either by: 1) retention or transformation of a capillary present early in embryonic or postnatal development (pre-existing connections) - arterialization (Mac Gabhann and Peirce, 2010), 2) *de novo* formation of collaterals by arteriolar sprouting – arteriogenesis (suggested by Lucitti et al., 2012); or 3) migration of arterial ECs from arteries along existing capillaries and reassembly into collaterals - artery reassembly (Das et al., 2019).

Previous studies (Chalothorn and Faber, 2010; Lucitti et al., 2012) suggest that in the mouse embryo, pial collaterals form as sprout-like extensions of arteriolar ECs, which reside on top of the underlying plexus. Although this process is considered to be dependent on VEGFR2-signaling, no formal proof of sprouting angiogenesis was provided (Lucitti et al., 2012). Furthermore, endothelial behavior at the cellular and subcellular (filopodia) levels was not resolved in these studies. In addition, previous studies differ in the developmental time point when collaterals are first observed: E14.5 (Lucitti et al., 2012) or E15.5 (Chalothorn and Faber, 2010). To date, no precise cellular mechanism of pial collateral development has been described.

Research into collateral regeneration after stroke has recently gained considerable attention. After distal middle cerebral artery occlusion (dMCAO) pial collaterals rapidly expand in a process termed outward remodeling. The dMCAO stroke model is deemed most suitable for a detailed examination of collateral outward remodeling, with one study indicating that the collateral diameter observed on day 7 following dMCAO persisted until day 65 (Okyere et al., 2018). While Okyere et al (Okyere et al., 2018; Okyere et al., 2020) showed that this process involves endothelial proliferation, no study has quantified and traced the lineage of proliferating ECs over the course of collateral remodeling.

To gain deeper insights into the mechanisms of pial collateral formation and remodeling, we investigated the developmental origin of collateral ECs by combining lineage tracing of arterial and microvascular ECs with whole mount pial surface imaging. We found that mature pial collaterals are mostly derived from an embryonic arterial lineage (87%) and, to a lesser degree, from an embryonic microvascular lineage (23%). We provide evidence for mosaic colonization of pre-existing capillaries by arterial and microvascular cells to be the process driving pial collateral formation. We also identify that collateral outward remodeling is achieved by local proliferation of arterial cells, rather than recruitment of the venous and microvascular lineage. Our data support a model where pial vasculature deploys different cellular mechanisms in development and regeneration (after stroke). In development, collateral “loops” form by immigration of artery- and plexus-derived ECs into existing capillary structures, whereas in regeneration, collateral remodeling relies on local EC self-replenishment by proliferation.

## Results

### Pial collaterals form as lumenized vascular segments between E14.5 and P21

It has previously been described that pial collaterals form in two phases: 1) growth phase – involving an increase in collateral formation (starting at E14.5/E15.5 and peaking between E16.5 and E18.5), and 2) postnatal phase – involving vascular remodeling (Chalothorn and Faber, 2010; Kalos and Theus, 2022; Perovic et al., 2022). However, thus far, collateral growth and remodeling/regression have been predominantly assessed by vessel counting at stereomicroscopic resolution.

To analyze pial collateral development at cellular and subcellular resolution, we visualized the pial vasculature between E12.5 and P21 using ICAM2 – to indicate lumenization (Stenzel et al., 2011, Franco et al., 2015), aSMA – to label smooth muscle cell progenitors and smooth muscle cells (Jung et al., 2017) and CD93 – to label all of the developing endothelium (Lugano et al., 2018). At E12.5, the middle cerebral artery (MCA) trunk and main branches are clearly identifiable (ICAM2+, CD93+) in contrast to the underlying plexus (only CD93+). aSMA covers the MCA trunk, indicating smooth muscle cell progenitor recruitment as early as E12.5 (Figure 1A).

**Figure 1:**
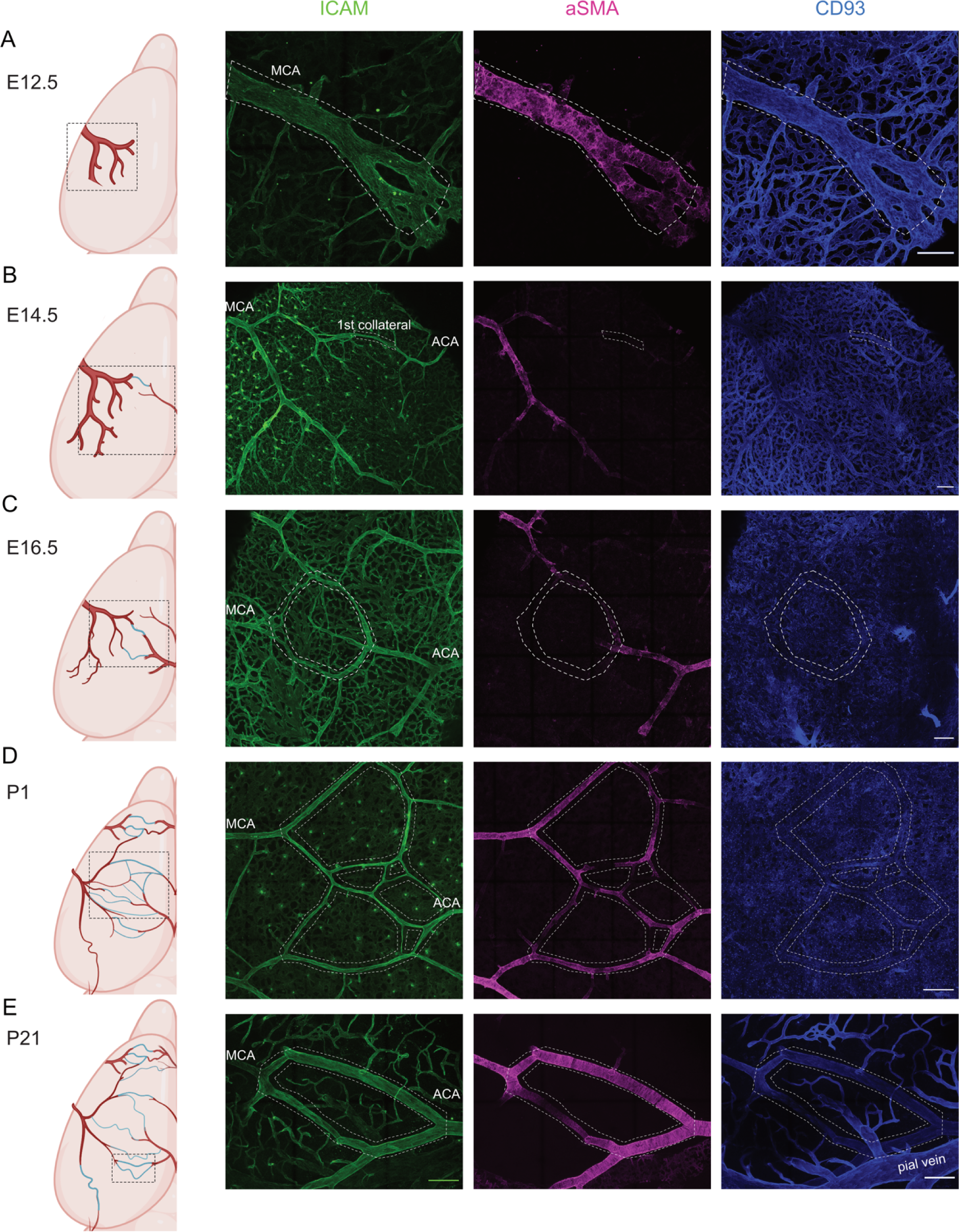
Pial collaterals form between E14.5 and P21, have a continuous lumen with arteries and recruit mural cells postnatally. Immunofluorescence staining against ICAM2 (green), aSMA (magenta) and CD93 (blue) was performed throughout the pial arterial development. Scale bar: 100μm. A. At E12.5, the MCA trunk and main branches are lumenized and show mural cell progenitor recruitment. White dotted lines indicate the main trunk of nascent MCA. B. First collateral vessels (white dotted square) are observed on the pial surface at E14.5, appear lumenized (ICAM2) arching over a dense venous plexus (CD93) and are devoid of aSMA coverage. C. At E16.5, pial collateral network is more complex, still intimately connected to the underlying plexus (now also ICAM2 positive) and still aSMA-negative. White dotted lines indicate an example of a collateral loop. D. At P1, pial collateral network is clearly a part of the superficial arterial circulation (ICAM2), shows a more complete aSMA-coverage and lies on top of a very dense venous plexus. White dotted lines indicate collateral network at P1. E. At P21, mature pial collaterals display an even more complete aSMA-coverage and are now surrounded by well-defined veins, venules and the residues of the pial plexus (CD93). White dotted lines indicate mature collaterals.

Between E14.5 and E18.5, there were many collateral loop-like connections between neighboring distal-end arterioles of different arterial trees (middle, anterior, and posterior cerebral arteries - MCA, ACA and PCA). We observed the first collateral connections between the MCA and ACA at E14.5 (Fig. 1B). These connections are prominently ICAM2 positive, whereas the underlying plexus is only faintly ICAM2 positive. Between E14.5 and P1, CD93 labels the dense plexus. By E16.5 (Fig. 1C), ICAM2 highlighted formation of more collateral connections, which increased manifold from E15.5 to E18.5 (as previously described by Chalothorn & Faber, 2010). At 16.5 (Fig. 1C), the entire arterial circulation is ICAM2-positive, whereas aSMA-labeling is only present in the bigger branches of MCA and ACA, and is still absent in collaterals. By P1 (Fig. 1D), the arterial network shows more complete aSMA coverage. At this point in development, there is a dense, faintly ICAM2-positive plexus under the arterial circulation, which is mostly pruned out by P21 (Fig. 1E). This is in agreement with previous studies (Letourneur et al., 2014; Coelho-Santos and Shih, 2020) on postnatal pruning of the superficial plexus. By P21, the collateral network is still ICAM2-positive and shows more complete aSMA labeling. Together, these observations suggest that pial collaterals first become visible as part of the pial arterial circulation at E14.5 and recruit smooth muscle progenitor cells from P1 onwards.

### Mature collaterals are mosaic in origin

To understand and quantify the contribution of arterial or microvascular ECs to forming collaterals, we used two Cre-Lox inducible mouse lines as lineage tracing tools: Bmx-CreERT2 (Ehling et al., 2013) for arteries and Vegfr3-CreERT2 (Martinez-Corral et al., 2016) for microvasculature/veins, in combination with the R26mTmG reporter line (Muzumdar et al., 2007).

We induced tamoxifen-mediated reporter labeling at E13.5, 24h before we observed the first collaterals forming, in order to genetically and irreversibly label by GFP the Bmx- or Vegfr3-expressing ECs of the nascent pial arterial circulation or pial capillary plexus. These embryos expressed GFP in arterial or capillary ECs upon tamoxifen injection (labeling frequency, 92% of all arterial ECs are labeled with Bmx-CreERT and 6,2% of all microvascular ECs are labeled for Vegfr3-CreERT) (Fig. S1A,B, F, and H). We determined the time window in which a single tamoxifen injection leads to recombination. This is essential to be able to distinguish between *bona fide* lineage tracing, and potential later recombination events during the peak formation of collaterals. CreERT protein localization was previously used to determine the time window of tamoxifen activation after a single injection (Hayashi & McMahon, 2002; Cano et al., 2023). The CreERT fusion protein (Cre-recombinase fused to a fragment of the Estrogen receptor alpha, ERα) relocates to the nucleus during the activation time window and relocates back to the cytoplasm at the end of the activation. We examined the subcellular localization of CreERT by immunostaining against the ERα protein domain. After the first 24 hours post-tamoxifen (at E14.5), CreERT was localized almost exclusively in the nuclei of Bmx-expressing cells (Fig. S2A). At E15.5, CreERT was localized both in the nucleus and cytosol of the Bmx-expressing cells (Fig. S2B). At E16.5, CreERT2 was mostly expressed in the cytosol, indicating CreERT2 translocation back to the cytosol (Fig. S2C). These results indicate that with our Tamoxifen dose, BmxCreERT is only active in the nucleus between E13.5 – E15.5, validating that any population we find labeled at later time points derives from ECs that expressed *BMX* between E13.5 and E15.5.

Our confocal imaging of the entire pial surface enabled us to anatomically distinguish collaterals from neighboring arterioles. Mature collaterals are easily identifiable in the adult as the often tortuous connections between the neighboring arterioles of the highest branching order (Faber et al., 2014; Kaloss & Theus, 2022; Perovic et al., 2022). In the embryo, collateral connections show no tortuosity and resemble vascular loops (Fig. 1C).

Our analysis of the BmxCreERT;R26mTmG line showed that at E18, 74% of all collateral ECs were derived from Bmx-lineage (Fig. 2A,E). In contrast, only 18% of all ECs in collaterals were of Vegfr3-lineage (Fig. 2B,E). Next, we traced the Bmx- and Vegfr3-embryonic lineages until P21. Similarly, at P21, 87% of all collateral ECs were derived from the embryonic Bmx-lineage (Figure 2C,F); only 23% of cells in mature collaterals are derived from the embryonic Vegfr3-lineage (Fig. 2D,F). Collectively, these lineage tracing results indicate that embryonically labeled arterial cells between E13.5 – E15.5 contribute to most collateral vessel formation, both in late embryogenesis and the postnatal mature state.

**Figure 2:**
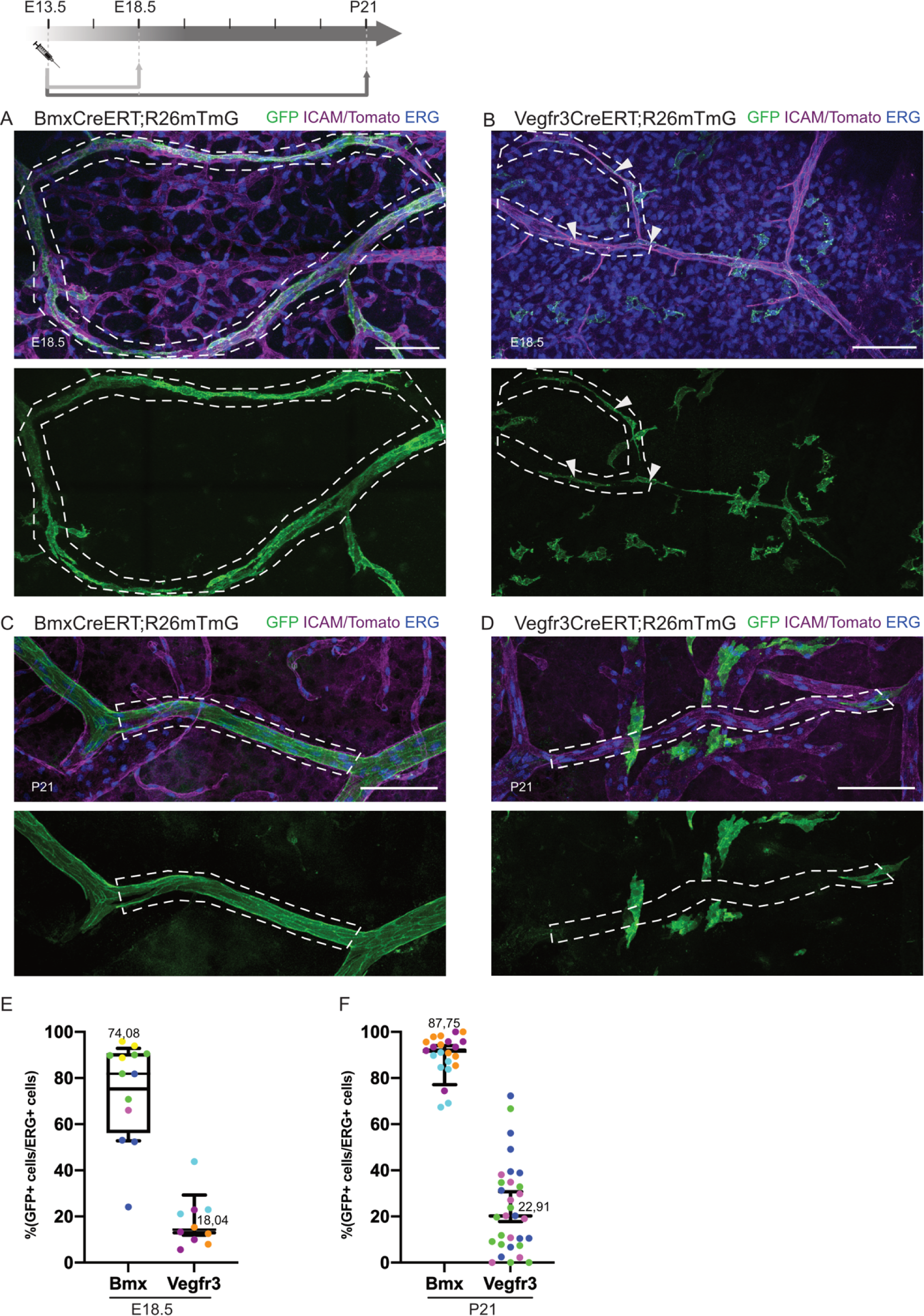
Mature collaterals are mosaic in origin: composed of mostly Bmx-(arterial) and, to a lesser degree, Vegfr3-(microvascular) endothelial lineage. Pregnant dams were injected with tamoxifen at E13.5: samples were harvested at E18.5 (A,B) or P21 (C,D), as indicated in the experimental scheme (top left) and immunolabeled against ICAM2, GFP and endothelial nuclear marker, ERG. White dotted lines represent collateral loops (A,B) or single mature collaterals (C,D). Scale bars: 100um. GFP (lineages) are shown separately under each merged image. A. E18.5 BmxCreERT;R26mTmG pial collaterals. Bmx-lineage cells (GFP) label the majority of the nascent collateral. B. E18.5 Vegfr3CreERT;R26mTmG pial collaterals. Some of the Vegfr3-lineage ECs (GFP) are embedded in the forming collaterals and in contact with the lumen (indicated by white arrows), while most of them belong to the plexus. C. P21 BmxCreERT;R26mTmG pial collateral. Embryonic (E13.5) Bmx-lineage labeling of the mature collateral. D. P21 Vegfr3CreERT;R26mTmG pial collateral. Embryonic Vegfr3-lineage labels some cells in mature collaterals. E. Percentage of Bmx- or Vegfr3-lineage, quantified as the number of double positive GFP+ERG+ cells over all ERG+ cells within embryonic collaterals (E18.5) F. Percentage of Bmx- or Vegfr3-lineage, quantified as the number of double positive GFP+ERG+ cells over all ERG+ cells within mature collaterals (P21). N≥3 animals. Mean and SD are calculated using average data points per animal. For transparency reasons, all data points are represented as scatter dot plots and individual animals are color-coded.

### Postnatal growth of pial collaterals predominantly involves Bmx-lineage

Although pial collateral numbers decrease between E18.5 and P21 (Chalothorn and Faber, 2010), the postnatal cortical surface extends multifold (Kobayashi, 1963; Coelho-Santos and Shih, 2020), suggesting that some collateral vessels regress and others grow in length and potentially diameter, incorporating new cells. In order to understand whether the postnatal growth of the maintained collaterals relies on arterial or microvascular EC recruitment, we traced arterial circulation and microvasculature between P1 and P21. Here, it is important to note that P1-labeling of BmxCreERT;R26mTmG line results in very high labeling frequency of P2 arteries and collaterals (97,9%); whereas P1-labeling of Vegfr3CreERT;R26mTmG line results in labeling of 19,9% of all microvasculature at P2 and very few cells in arteries (Fig. S1C-H).

Our analysis of the postnatal trace shows that at P21, 80% of all ECs in collaterals are derived from the P1-labeled Bmx-cells (Fig. 3A,E). In contrast to the embryonic trace, only 7% of all ECs in mature collaterals are derived from the P1-labeled Vegfr3-cells (Fig. 3B,E). Together, these results indicate that the growth of the sustained collaterals in the postnatal period involves primarily the Bmx-derived (arterial) ECs and, to a much lesser extent than in the embryo, the Vegfr3-derived (microvasculature) ECs. These results suggest that postnatal growth of pial collaterals relies on self-replenishment of arterial ECs.

**Figure 3:**
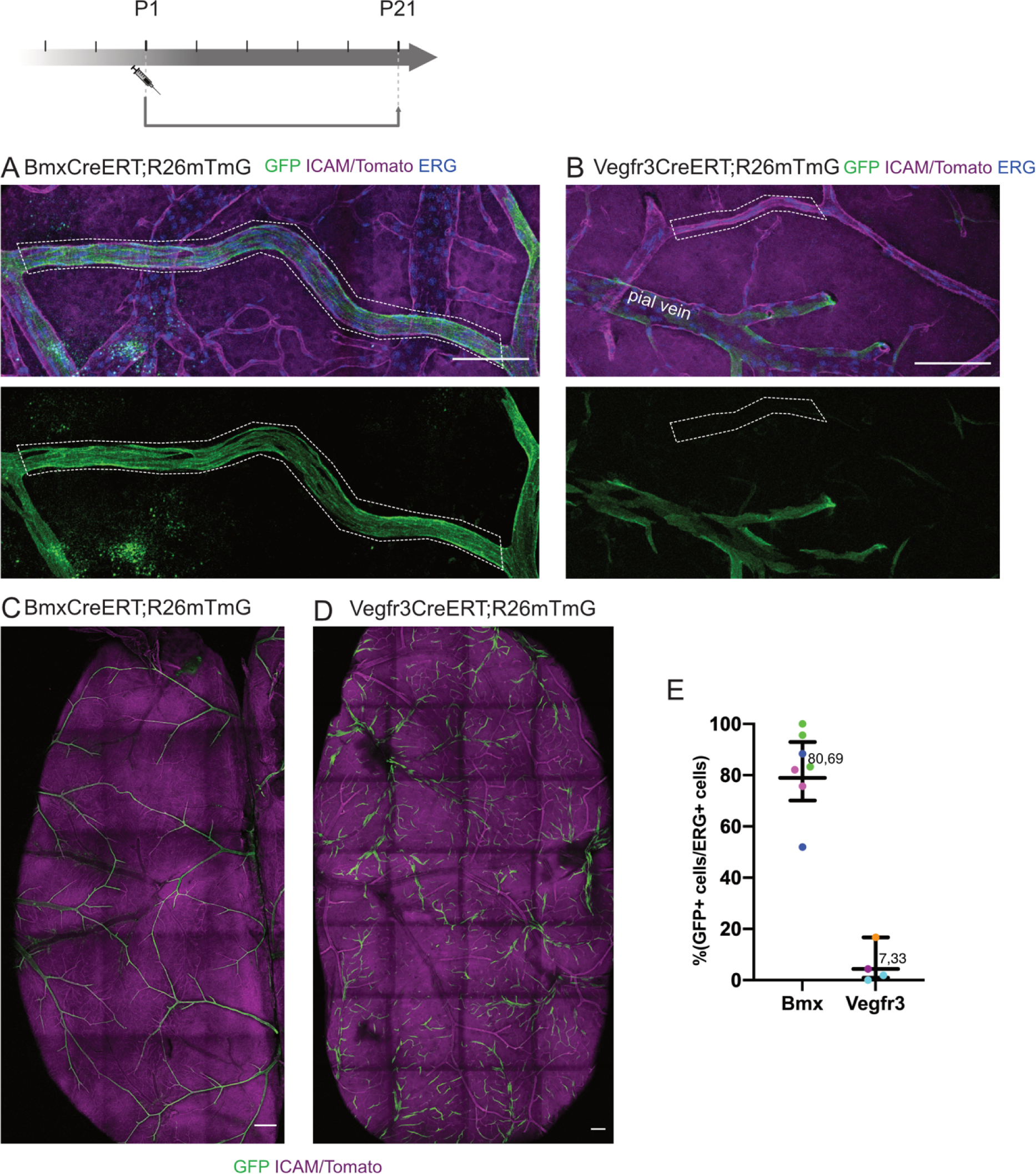
Postnatal growth of collaterals involves mostly Bmx-lineage. Pups were injected with tamoxifen at P1 and samples were collected at P21 and immune-stained against GFP (green), ICAM2 (magenta) and ERG (blue), as indicated in the experimental scheme (top left). White dotted lines represent individual collaterals (A,B). Postnatal Bmx-lineage (GFP) labels most of mature collateral vessels and arteries (A, C) whereas postnatal Vegfr3-lineage contributes much less to collaterals (B,D) and is mostly located around veins and venules at P21. Contribution of each lineage was quantified as the number of GFP+ERG+ within all ERG+ cells in mature collaterals and depicted in E. Percentage of Bmx- or Vegfr3-lineage, quantified as the number of double positive GFP+ERG+ cells over all ERG+ cells within mature collaterals. N=3 animals. Each data point represents one imaged collateral zone. Data are mean ±SD. Mean and SD are calculated using average data points per animal. For transparency reasons, all data points are represented as scatter dot plots and individual animals are color-coded.

### Collateral forming ECs display migratory phenotypes within already existing vessel structures

To understand the mechanism of pre-collateral formation in the embryo, we focused on stages E16.5 and E18.5. At E16.5, the pial network is still forming, and it shows a stronger ICAM2 labeling than the underlying plexus (Fig. 1C). Collaterals form at the intersection of already formed arteries and the plexus underneath. When studying in greater detail the zone between arterial trees, we noticed that there are vascular structures that connect to the feeding artery, often have a higher diameter than the plexus, show more pronounced ICAM2-signal and specifically recruit Bmx-lineage cells (Fig. 4A, S3A-D) and few Vegfr+ cells (Fig. 4B). The Bmx-lineage cells also exhibit migrational polarity away from the arterioles and towards the middle of these vessels, which is visible by the direction of their filopodia protrusions (observed as cellular protrusions of Bmx-lineage GFP+ cells). We termed these vessel structures *pre-collaterals*. At E18.5, pre-collateral vessels become even more pronounced against the background of the surrounding and underlying microvasculature and are similarly occupied by mostly Bmx lineage-derived ECs (Fig. 4C) and some Vegfr3 lineage-derived ECs (Fig. 4D). Again, we observed that the GFP+ cells of both lineages show a highly exploratory and migratory phenotype (Fig. 4C and D detail). However, we have not observed arterial tip-cell sprouting behavior, in terms of *de novo* arteriogenesis where single arterial tip cells invade interstitial spaces to form new collaterals. Instead, we observe highly active, migratory cells with polarized morphology undergoing intercalation movements into existing *pre-collateral* vessels. We therefore derive that in the embryo, collaterals form in a process of mosaic colonization of *pre-collateral* vessels (marked by pronounced ICAM2-signal and direct connection to arteries) by Bmx- and Vegfr3-lineage ECs.

**Figure 4:**
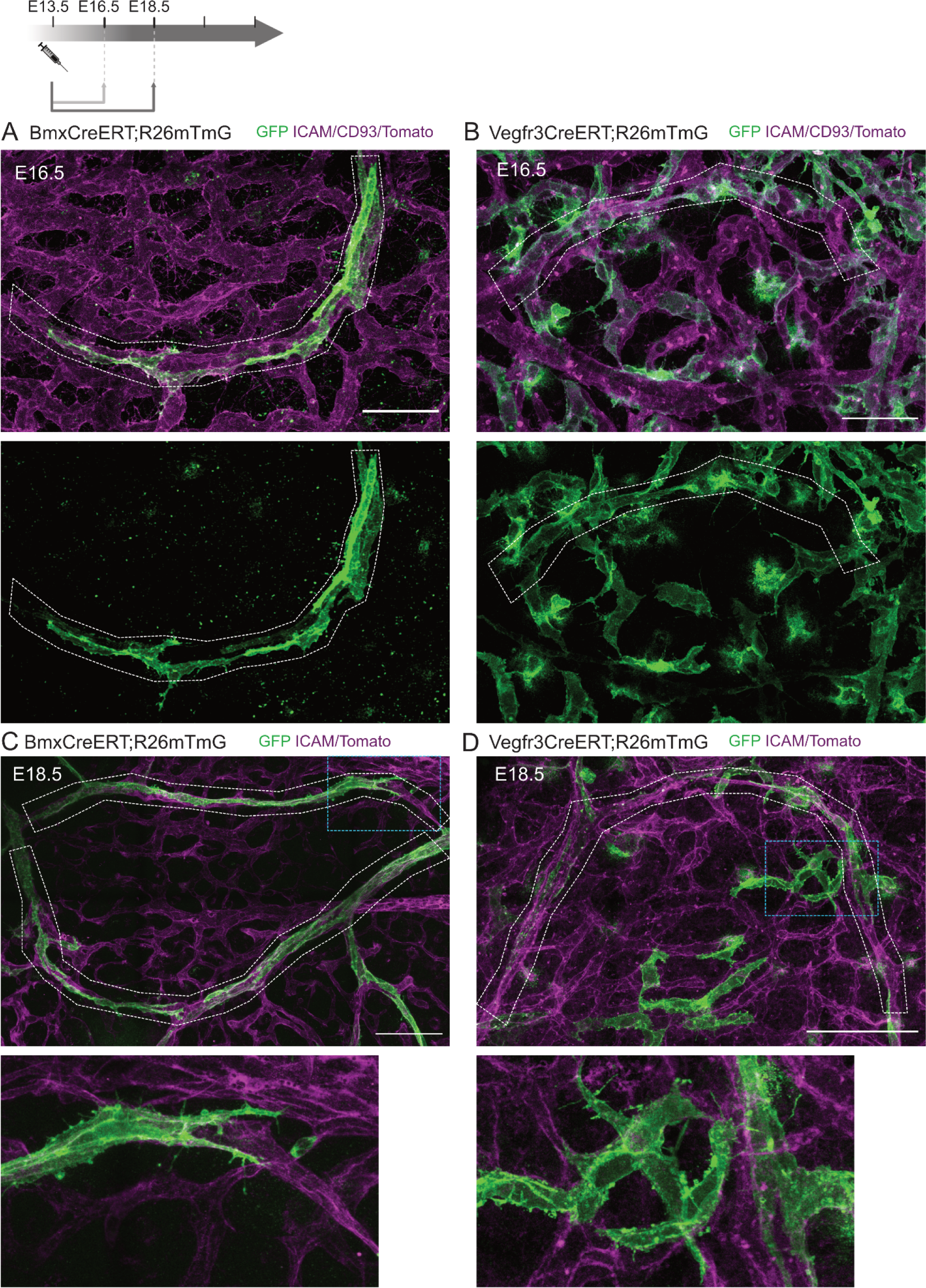
Bmx- and Vegfr3-lineage cells appear to be actively migrating within already existing capillaries. Bmx- and Vegfr3-lineage ECs were traced from E13.5 - E16.5 and E13.5 - E18.5, as shown in the experimental plan (upper left). GFP (green), ICAM2/CD93/Tomato (magenta). GFP-channel is shown separately A and B, respectively. White dotted lines represent each individual pre-collateral A. E16.5 BmxCreERT;R26mTmG pial pre-collaterals. B. E16.5 Vegfr3CreERT;R26mTmG pial pre-collaterals. C. E18.5 BmxCreERT;R26mTmG pial pre-collaterals. D. E18.5 Vegfr3CreERT;R26mTmG pial pre-collaterals. Blue dotted lines indicate vascular segments that are enlarged in the lower panel of C, D.

### Mosaic colonization of pre-collateral vessels coincides with their arterialization

Connexin 40 (Cx40) is one of the gap junction proteins which is highly expressed in mature arteries (Little & Beyer, 1995; Buschmann et al., 2010; Su et al., 2018; Cui et al., 2015; Márquez et al., 2023; Cano et al., 2023). Cx40 expression is upregulated in the context of high shear stress (Denis et al., 2019). Given the role of Cx40 in artery maturation and higher flow, we closely analyzed the relationship between Bmx-/Vegfr3-lineage ECs and Cx40 immunolabeling signal. As described above (Fig. 4), pre-collateral vessel connections at E16.5 contain GFP+ cells of predominantly Bmx-lineage and, to a smaller degree, of Vegfr3-lineage (Fig. 5A, B). We found that at E16.5, Cx40 expression is highest in the most established, wider arteries and arterioles (Fig. 5C, D, S4B, C), suggesting Cx40 to be a true marker of arterialization in pial circulation. Moreover, plexus structures that appear wider, and are judged as pre-collaterals show Cx40 expression in a pattern consistent with Cx40 being induced selectively in vessels of higher flow. We observed that Cx40 marks pre-collaterals both with and without already intercalated GFP+ cells (white arrows indicate Cx40+ GFP+ pre-collateral vessels in Fig. 5C-F). Intriguingly, sometimes even cells of Bmx-lineage were Cx40-negative, suggesting that commitment of lineage-positive cells to form arterial structures and their maturation are distinct and uncoupled events. The intercalation movements driving mosaic colonization and cell displacements can precede or coincide with the maturation of pre-collaterals marked by acquisition of Cx40 expression that is likely driven by increasing flow. Accordingly, smaller caliber vessels with Bmx-lineage ECs showed only faint Cx40-staining (white arrows in Fig. 5A, C). Likewise, smaller vessel segments with Vegfr3-lineage cells showed no Cx40 signal (yellow arrow in Fig. 5B,D,F). Together, these results suggest that migration of chiefly Bmx- and some Vegfr3-lineage ECs into forming collaterals happens concurrently with - but uncoupled from - the arterialization of pre-collateral vessels.

**Figure 5:**
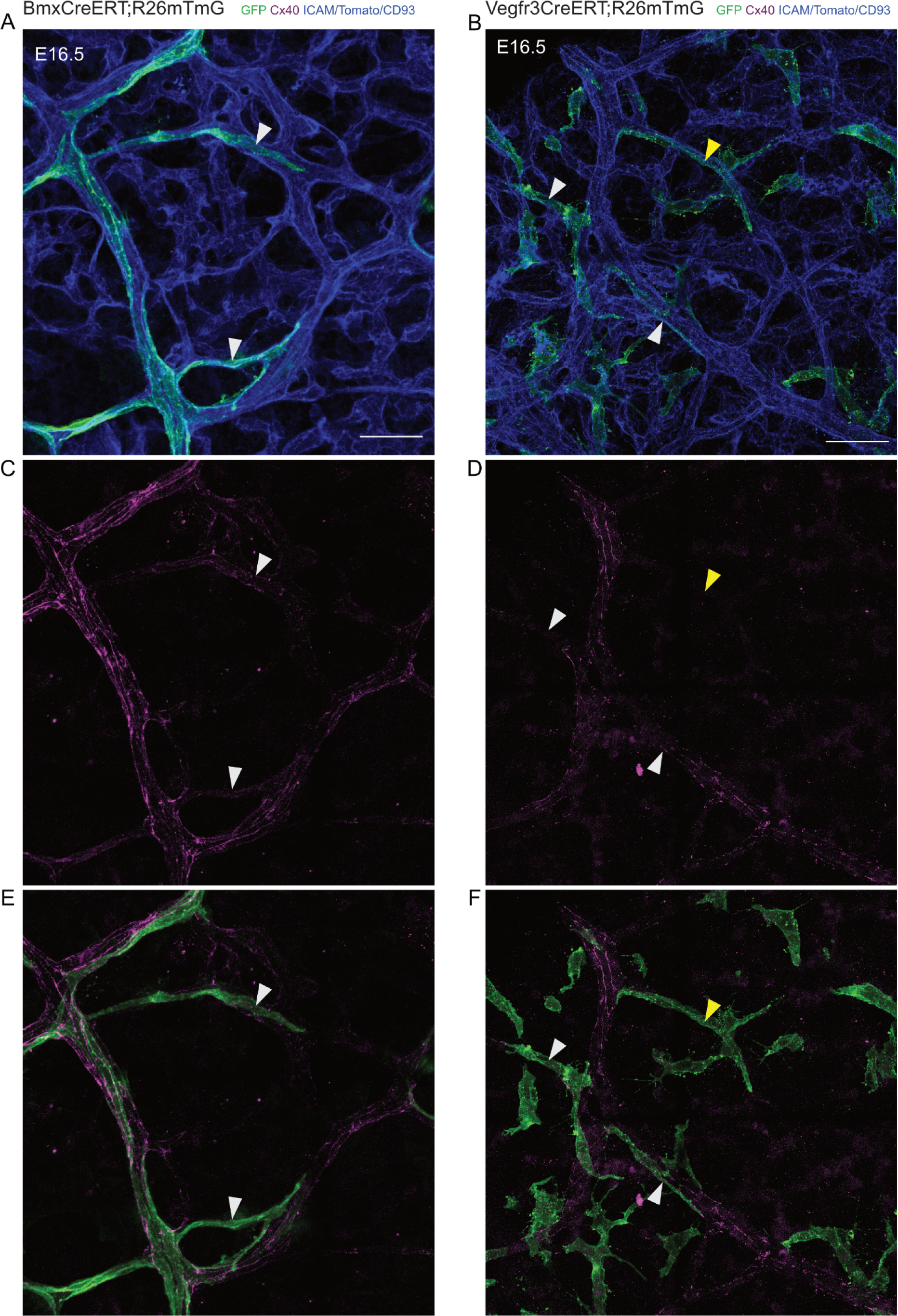
Bmx- and Vegfr3-lineage ECs mark pre-collateral vessels which are undergoing arterialization. Bmx- and Vegfr3-lineage ECs were traced from E13.5 - E16.5. GFP (green), Cx40 (magenta), ICAM2/CD93/Tomato (blue). ECs derived from Bmx-lineage were found in wider arterial segments which showed Cx40-positive staining and in narrower pre-collateral segments (faintly Cx40-positive) - indicated by white arrowheads (A,C,E). ECs derived from Vegfr3-lineage were found both in vessel segments with weak Cx40 staining (white arrowhead) and Cx40-negative vessel segments (yellow arrowhead) (B,D,E). N=4 animals from two di?erent litters.

### Arterial cells proliferate during collateral formation

Arterial ECs rarely proliferate, and cell cycle exit appears to be even required for arterial cell specification (Su et al., 2018; Luo et al., 2021). If pial arteries were to stop proliferating, where would the pre-collateral invading, Bmx-lineage ECs come from? To answer this, we performed EdU injections into pregnant females. We assessed pial EC proliferation at E16.5 and E18.5, at times of peak collateral formation. ICAM2 staining and vessel morphology was used to distinguish between embryonic arteries and the surrounding, underlying plexus. Proliferating ECs (in S phase) were quantified by counting double-positive cells for EdU and ERG. Our analysis shows that at E16.5, 3% of pial arterial ECs proliferate (Fig. 6A,C); at E18.5, 4% of pial arterial ECs proliferate (Fig. 6B,C). In contrast, at E16.5 14% of all pial ECs proliferate (Fig. 6A,C) and at E18.5 12% of all pial ECs proliferate (Fig. 6B,C). Thus, we conclude that embryonic pial arterial ECs do proliferate, although they do so to a lesser degree than EC in the underlying pial plexus.

**Figure 6:**
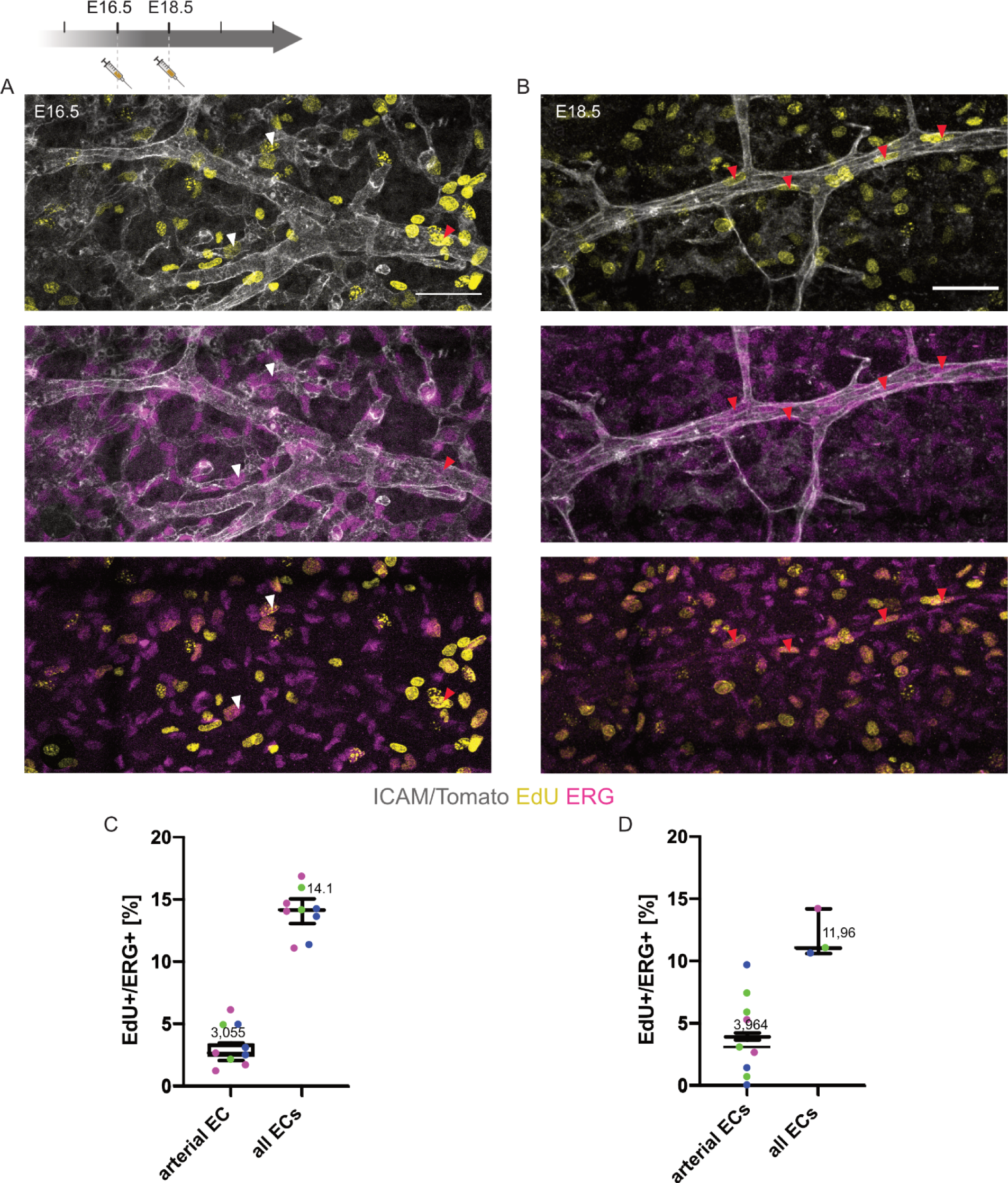
Arterial cells proliferate during collateral formation. EdU injections were performed at E16.5 and E18.5, 2h before sacrifice. Upper panel shows ICAM/Tomato (gray) and EdU (yellow) staining, middle panel shows ICAM/Tomato (gray) and ERG (magenta) staining and lower panel shows ERG (magenta) and EdU (yellow) labeling. At E16.5 there were double positive cells (EdU+ERG+) in pial arteries and the surrounding plexus (A). At E18.5, we also found proliferating endothelial cells (EdU+ ERG+) in both arteries and plexus. Red arrowheads indicate examples of double positive (ERG+EdU+) arterial cells. C. Quantification of proliferating ECs at E16.5. D. Quantification of proliferating ECs at E18.5. N=3 animals per stage. Mean and SD are calculated using average data points per animal. For transparency reasons, all data points are represented as scatter dot plots and individual animals are color-coded.

### Outward remodeling of collaterals after stroke (dMCAO) relies on local proliferation of arterial endothelial cells

Pial collaterals have a unique ability to undergo dramatic outward remodeling in the event of stroke (Coyle et al., 1991; Wei et al., 2001). Collateral capacity to expand the vessel diameter relies on cell proliferation (Okyere et al., 2016). However, the source of proliferating cells which integrate into the growing collateral has not been identified. Given the high proliferative capacity of venous ECs (Xu et al., 2014; Lee et al., 2021), we tested the hypothesis that in stroke, veins and the microvasculature might represent the main source of proliferating cells in growing collaterals. To do so, we combined our postnatal lineage tracing with distal middle cerebral artery occlusion (dMCAO), followed by daily EdU injections for a week. Our analysis showed that Bmx-lineage contributed to 96% - 100% of all ECs in collaterals of both sham and ipsilateral (stroke) animals and upstream MCA branches (Fig. S5), which is an even higher number than in our postnatal lineage trace (81% P1 Bmx-lineage contribution to P21, Fig. 3E). Next, we observed that ipsilateral collaterals exhibit a bulging morphology signifying diameter increase (Fig. 7A,D). 20% of all ECs in the remodeled collaterals were EdU+, and all of them were Bmx-lineage derived (Fig. 7A,G). In contrast, we found no cells of Vegfr3-lineage in remodeling collaterals (Fig. 7D). Collaterals in sham operated animals and collaterals of the contralateral hemisphere exhibited the tortuous morphology characteristic of mature quiescent collaterals with no observed bulging (Fig. 7B,C). In collaterals of the sham control, we found only 3% of cells to be EdU+ (Fig. 7B,G). Intriguingly, 18% of ECs within the ipsilateral upstream MCA branches of the remodeled collaterals were proliferating (Fig. 7E, H). In contrast, the sham arteries had very few proliferating ECs (0,3%) (Fig. 7F, H). In summary, these results indicate that one week after dMCAO, collateral remodeling relies on local proliferation of Bmx-derived ECs and does not involve recruitment of Vegfr3-lineage.

**Figure 7:**
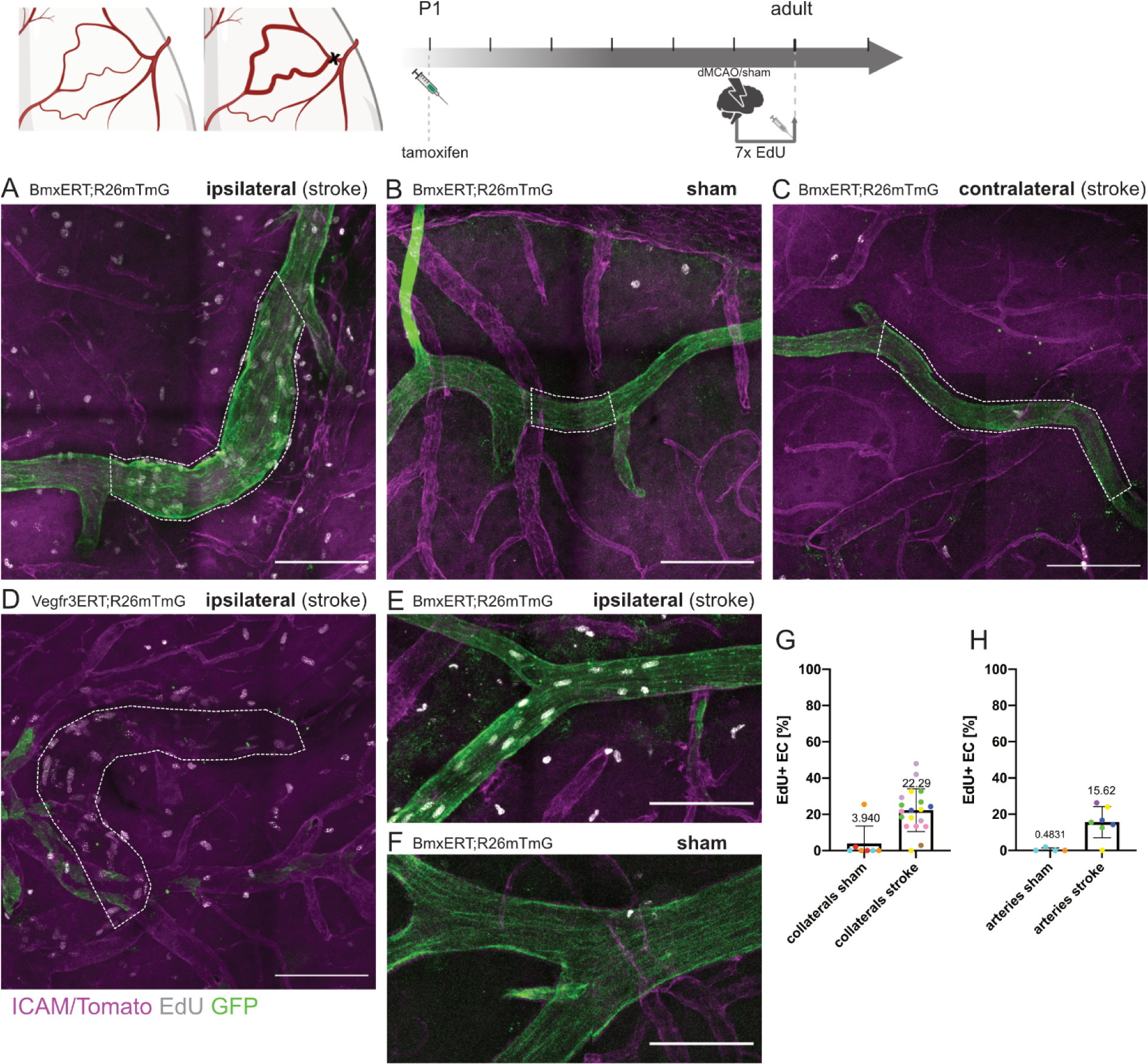
After stroke (dMCAO), collateral outward remodeling happens through proliferation of collateral (local) and nearby arterial endothelial cells and not through recruitment of microvascular cells. To combine lineage tracing and dMCAO, adult (8 - 12 weeks old) mice lineage-labeled at P1 underwent a dMCAO/sham procedure. Following operations, animals were injected with EdU daily for the next 7 days and then sacrificed. GFP (green), ICAM/Tomato (magenta) and EdU (gray). White dotted lines represent each individual collateral. Collaterals in animals with dMCAO underwent a massive remodeling, marked by their bulging morphology and many proliferating ECs in both lines (A,D). Collaterals from the sham BmxCreERT;R26mTmG animals or contralateral hemisphere of dMCAO animal group displayed normal morphology, GFP and few proliferating ECs (B,C). MCA branches directly upstream of the remodeling collateral showed marked EC proliferation in the dMCAO BmxCreERT;R26mTmG animals (F) and no proliferation in the sham BmxCreERT;R26mTmG animals (E). G. Quantification of EdU+ endothelial cells in ipsilateral collateral vessels (collaterals stroke) and sham control collateral vessels (collaterals sham). H. Quantification of EdU+ endothelial cells in ipsilateral upstream MCA branches (arteries stroke) and sham upstream MCA branches (arteries sham). N=2 animals for sham (technical control) and N=5 for stroke animals. Mean and SD are calculated using average data points per animal. For transparency reasons, all data points are represented as scatter dot plots and individual animals are color-coded.

## Discussion

The cerebral vasculature bears very little room for redundancies, due to its compelling complexity and regional specialization. Pial collaterals are thus a rare gem, as they represent a back-up feeding strategy. After ischemic blocking of the main feeding artery, collaterals undergo extensive remodeling in a matter of days, all while maintaining vascular integrity and function (Faber et al., 2014; Okyere et al., 2018; Kaloss and Theus, 2022; Maguida and Shuaib, 2023). Although collateral enhancement in patients has substantial clinical potential as an acute stroke therapy, collateral therapeutics remain a relatively uncharted therapeutic strategy (Winship, 2015; Iwasawa et al., 2016). This is partly due to a gap in knowledge on the precise cellular and molecular mechanisms of collateral formation and remodeling. Here we show that in mice, the first pre-collateral connections are already lumenized, increasing in number between E14.5 and E18.5 and obtaining mural cell coverage from P1 onwards (Fig. 1B-E). Embryonic arterial EC lineage labeling (Bmx-CreERT) amounted to 74% of E18.5 collaterals (Fig. 2A,E) and 87% of mature collaterals (Fig. 2C,E), whereas embryonic microvascular lineage (Vegfr3-CreERT) contributed to 18% of E18.5 (Fig. 2B,E) and 23% of mature collaterals (Fig. 2D,E). Next, we showed that postnatal growth of collaterals involves mostly arterial lineage, with very little contribution by microvascular cells (Fig. 3). We also found evidence that at the time of collateral formation, 3–4% of ECs in neighboring arteries proliferate (Fig. 6). In sum, most of the ECs that comprise mature collaterals are derived from an early embryonic arterial endothelial pool, rather than from venous/microvascular endothelium, in contrast to what has recently been shown for arteries in other organs (Xu et al., 2014, Lee et al., 2021, Giese & Rosa et al., 2023).

Additionally, we found that between E16.5 and E18.5, arterial (Bmx-lineage) and microvascular (Vegfr3-lineage) ECs migrate into already existing pre-collateral vessels (Fig. 4). Based on the present observations, we propose that pial collateral vessels that form between E16.5 and E18.5 do so, not as arteriolar sprouts but in the process of migration of single cells of Bmx+ and Vegfr3+ lineages into already existing vessels. Similarly, lineage tracing of arteries in neonatal mouse heart collateral formation after injury showed that singular pre-existing arterial cells are recruited to the watershed region of an infarcted neonatal heart to form new collaterals (Das et al., 2019). We found Cx40 to be a true zonation marker of arterial maturation (Fig. S4) and often co-expressed with GFP in forming collaterals, suggesting that migration of artery- and microvasculature-derived ECs into these pre-collateral capillaries might trigger arterialization (Fig. 5). However, the fact that we observe Cx40 expression in structures that have not (yet) recruited Bmx-lineage ECs, and those that show Bmx-lineage ECs without Cx40 expression, demonstrates that the two events, commitment and recruitment of Bmx-lineage ECs, and the establishment of an arterial gene signature, are distinct and likely uncoupled events. Given the known role of shear stress in Cx40 expression (Buschmann et al., 2010, Cui et al., 2015, Márquez et al., 2023) and of the Cxcl12-Cxcr4 signaling axis in pre-arterial cell recruitment (Das et al., 2019), at least in the heart, it is tempting to speculate that the colonization of pre-collaterals is primarily chemokine driven, whereas their maturation is primarily flow/shear stress induced.

Here we report that the pial collateral network developed through mechanisms distinct from those described to date. Instead of the suggested *de novo* arterial tip cell sprouting – arteriogenesis (Lucitti et al., 2012) or mechanisms found in other organs such as arterialization (Mac Gabhann and Peirce, 2010) or artery reassembly (Das et al., 2019), embryonic brains use a unique cellular process that is a combination of all three. It utilizes newly derived arterial and plexus cells (resembling arteriogenesis) and relies on migration of these arterial cells (resembling artery reassembly) into already existing capillaries (resembling arterialization). We termed this process of vessel formation mosaic colonization, as two distinct sources of cells are recruited to an existing vessel, thereby configuring a collateral. Overall, it is most similar to artery reassembly, except we found that there is a second, non-arterial source of endothelial cells in the forming pial collaterals, the Vegfr3-lineage. However, neo-collateral formation in adults may follow different principles.

Several reports have shown that pial collaterals require proliferation to remodel and expand their diameter in response to permanent middle cerebral artery occlusion (Okyere et al., 2018; Okyere et al., 2020). Our interest in collateral lineage analysis was partly sparked by the hypothesis that endothelial cells of different developmental origins respond differently to stressors (e.g., ischemia), as has been found for other lineages (Majesky, 2007). Using an exploratory study design, here we demonstrate that ∼ 20% of endothelial cells within a remodeling collateral proliferate within one week after stroke surgery (Fig. 7). Additionally, our analysis identifies that the vast majority (97%) of the cells in remodeling, ipsilateral collaterals are of arterial origin and that all of the proliferating cells can be traced from the arterial (Bmx) lineage we induced at P1 (Fig. 7, S5). In opposition to the 96 - 100% of P1-arterial labeling in ipsilateral/sham collaterals, P1-arterial trace results in 80% labeling of P21 collaterals (Figure 3A,C,E) and P1-arterial trace results in 89% labeling of adult contralateral collaterals of stroke animals (Figure 7C, 5S). Of note, animals undergoing dMCAO/sham were on average 2-3 months old and we did not analyze the fate of Bmx/Vegfr3-lineage ECs in collaterals between P21 and 2-3 months mice. We can only speculate that Bmx-lineage ECs outweigh Vegfr3-lineage between P21 and 2-3 months of animals’ age. Since we observed a moderate increase in the Bmx-derived EC fraction on the contralateral side from 80% to 89%, but a steeper increase in Bmx-derived EC fraction on the ipsilateral/sham side 96-100%, it is interesting to speculate that the injury itself prompts Bmx-derived ECs to outcompete Vegfr3-derived ECs in ipsilateral collaterals. In toto, these data suggest that collateral remodeling at least partly depends on proliferation of local artery-derived cells (within the growing collateral and in the MCA - directly upstream). Our findings demonstrate a surprising proliferative capacity of arterial cells, normally believed to be difficult to reactivate.

The last decade established that endothelial cells generally migrate from low flow segments towards higher flow segments during remodeling. In the mouse yolk sack and zebrafish brain vasculature (Udan et al., 2013; Chen et al., 2012), vessel segments with lower flow levels undergo regression. In the mouse retina, this flow-migration coupling leads to pruning of the plexus (Franco et al., 2015) and immigration of venous ECs into arteries (Lee et al., 2021; Giese & Rosa et al., 2023). Surprisingly our data demonstrate that collateral formation follows different rules, as artery-derived ECs migrate within the plexus in a fashion that contradicts migration against the flow direction. Taken together, our findings that the majority of mature collateral cells derive from embryonic arterial lineages and that these intercalating artery-derived cells in pre-collaterals show migrational polarity away from the arterioles suggest that arterial cells migrate with the flow, even though there is only very low flow at this point, as predicted based on previous embryonic flow assessments (Jones et al., 2004; Buschmann et al., 2010). Our data raises the question of what are the drivers of migration that attract the ECs of Bmx- and Vegfr3-lineages towards pre-collaterals. Moreover, what is the true significance of the Vegfr3-lineage contribution? Are the lineages mutually complementary or redundant? Increasing evidence suggests that neonatal coronary collateral formation is guided by the CXCR-4-CXCL12 signaling axis, as well as the Vegfr2 signaling network (Das et al., 2019, Arolkar et al., 2023). During neonatal collateral artery reassembly, CXCR-4 (receptor) is expressed by pre-existing arterial cells, whereas CXCL12 (ligand) is secreted by microvasculature, thus serving as a chemical guide for nascent collateral formation. Endothelial-specific knockdown of *Cxcr4* in adult mice abolishes hypoxia-driven pial neo-collateral formation (Zhang et al., 2020). Whether CXCR-4 - CXCL12 also guides embryonic pial collateral formation will require further work.

Furthermore, our findings invite the question of whether collateral mosaic origin is indicative of continuous heterogeneity. We find that at P2, the vast majority of collateral ECs express Bmx (Fig. S1, C-H), suggesting that the embryonically derived Vegfr3+ collateral cells lose their Vegfr3-expression and start upregulating *BMX*. This indicates that the endothelial plasticity we find in the embryo, where microvascular cells help build collaterals, might be lost postnatally. However, we still don’t know whether cells coming from the embryonic Vegfr3-lineage carry molecular memory. Finally, are the collateral ECs of arterial and microvascular lineages functionally distinct? We described that, upon stroke, postnatal Bmx-lineage takes center stage in expanding the collateral vessel. Is having once Vegfr3-derived, but now Bmx+ cells in mature collaterals somehow advantageous to having all cells in collaterals be derived from Bmx-lineage and remain Bmx+ throughout their lives? This connection between origin and function in collateral biology remains to be further explored.

Taken together, we propose a novel, integrated model of embryonic pial collateral formation (Fig. S6), based on the first ever lineage tracing study of pial collaterals. We termed it *mosaic colonization*, implying that arterially and capillary-sourced ECs are recruited into an already existing vessel structure (pre-collateral), converging with arterialization of this pre-collateral segment. We find that collateral outward remodeling, in contrast, only involves proliferation of local artery-derived ECs within the already existing collateral, suggesting that the developmental program of mosaic colonization is not reactivated during adaptation to injury. Our exploratory findings provide mechanistic insight into how collaterals form and grow and will hopefully set the stage for further development of ways to pharmacologically condition or reactivate collaterals in stroke patients.

## Materials and methods

### Mouse lines

All mouse lines used have the same C57/BL6J background. The following mouse strains were used: BmxCreERT (Tg(Bmx-cre/ERT2)1Rha) (Ehling et al., 2013), Vegfr3CreERT (Tg(Flt4-icre/ERT2)Sgo) (Martinez-Corral et al, 2016) and R26mTmG (Gt(ROSA)26Sortm4(ACTB-tdTomato,-EGFP)Luo/J) (Muzumdar et al., 2007). Mice were maintained at the Max Delbrück Center for Molecular Medicine under standard husbandry conditions and were handled in compliance with the institutional and European Union guidelines for animal care and welfare. All embryos were staged from the time point of vaginal plug, which was designated as embryonic day 0 (E0). Single injections of tamoxifen (Sigma-Aldrich) were performed intraperitoneally (20 µg per g of animal). Pregnant dams were intraperitoneally injected with tamoxifen at embryonic day 13.5 To induce Cre-mediated recombination in postnatal day 1 (P1) pups, 4-hydroxytamoxifen (Sigma, 7904) was injected intraperitoneally (IP) (20 μL/g of 1 mg/mL solution) at postnatal day 1 (P1). Male and female mice were used for the analysis. Animal procedures were performed in accordance with the animal license G 0204/19.

### Stroke model (distal MCAO) with craniotomy

Anesthesia is induced by isoflurane 2% in 70% N2O and 30% O2 or maintained by ketamine/xylazine i.p. for the duration of the procedure. The gas mixture is delivered to the spontaneously breathing animals via a mask. Body temperature is maintained between 37 - 37.5 °C by a rectal temperature probe and a temperature-controlled heat mat. The skin area between the left eye and ear is disinfected. Before beginning the surgical procedure, the animal’s paw is pinched with forceps to check for the absence of pedal reflex (indicator of proper anesthesia). Eye ointment is applied to the eyes at the beginning of the experiment to protect them against dehydration. Lidocaine 0.5% is injected subcutaneously in the area of the skin incision before the start of the surgical procedure (preemptive analgesia, the total dose does not exceed 7 mg/kg). A skin incision is made centrally between the eye and the ear in a dorso-ventral direction. To extend it to a T-shaped incision, it is widened at the upper edge in caudo-rostral direction by about 4 mm each. The left temporal muscle is cauterized at its apical and distal parts, then detached from the skin and carefully pushed aside without injuring the retro-orbital vein. The wound is approximately 0.8 cm in length. The temporal muscle is knocked aside and fixed (e.g., with a suture or skin retractors) to expose the skull over the MCA territory. Using a drill, the bone is thinned (diameter approx. 2 mm) and then carefully lifted and removed with fine forceps. NaCl solution is applied between the drilling steps to avoid overheating due to the drilling. After identification of the distal middle cerebral artery, an electrocoagulation forceps is used to coagulate the vessels. Sham animals underwent the same procedure except the electrocoagulation of the distal MCA.

### Immunofluorescence staining

Embryos and pup brains were fixed in 4% fresh paraformaldehyde (PFA) solution in PBS overnight rocking at 4°C and washed in PBS. All brains were permeabilized overnight with TNBT (0.5% Perkin Elmer blocking reagent/0.05%Triton-X/3% 5M NaCl/10%Tris buffer pH 7.4/PBS) at 4°C. The following antibodies were used: anti-ICAM2 (BD Pharmigen, 553326), anti-GFP (Abcam, ab13970), anti-ERG (Abcam, ab92513), anti-CD93 (R&D Systems, AF1696), anti-Cx40 (Alpha Diagnostic, CX40-A), anti-aSMA-Cy3 (Sigma, C6198). Alexa fluor-conjugated secondary antibodies (Invitrogen) were used. Both primary and secondary antibodies were applied overnight at 4°C in TNBT buffer (dilution factor for primary antibodies is 1:100 and for secondary antibodies 1:200). Samples were washed with TNT buffer (0.05%Triton-X/3%5M NaCl/10%Tris buffer pH 7.4/PBS) at room temperature for a minimum of 6 hours (washing cycles between 30min and 1h).

### Pial surface confocal imaging

For imaging, samples were placed in a drop of PBS on a glass-bottomed dish. Images were taken at room temperature using LSM 780 and LSM 980 inverted microscope (Zeiss) equipped with a Plan-Apochromat 20×/0.8 NA Ph2 objective, Plan-Apochromat 25x/0.8 LD LCI objective, Plan-Apochromat 40x/1.4 DIC objective. Z-stacks of a maximum of 350µm were acquired. After confocal images acquisition, Zen 3.4 Blue edition software (Zeiss) was used for stitching. ImageJ software was used as an image analysis tool.

### In vivo EdU labeling and EC proliferation detection

To detect EC proliferation in embryonic brains, 50mg/kg body weight of thymidine analog EdU (Invitrogen, A10044) was injected intraperitoneally into pregnant dams 2h before dissection. Embryonic brains were isolated for whole mount analysis. In mice that underwent stroke, the same concentration of EdU was injected daily for 7 consecutive days after stroke. EdU signals were detected with the Click-it EdU Alexa Fluor 647 Imaging Kit (Invitrogen, C10340). In brief, after all other primary and secondary antibody incubations, samples were washed according to the immunofluorescence staining procedure and then incubated with Click-iT EdU reaction cocktails for 40 min, followed by DAPI staining.

### Statistical approach

Due to this study’s descriptive nature and its exploratory design, we abstained from statistical tests. Values shown are mean and standard deviation was used as the dispersion measure for a given animal. A statistical method of sample size calculation was not used during study design. We used an average of 3 animals per experiment, from different litters (detailed number of animals used in figure legends). Individual animals were color-coded.

### Methods to prevent bias

In our stroke experiments, all mice underwent MRI imaging at 24 h after the stroke. Animals with bleedings or extraterritorial ischemia or technical failures to occlude the dMCA were excluded. Sham mice with bleedings or massive edema were excluded. For technical reasons (identification of the collaterals and localisation at the ischemic cortex), blinding against animal groups was not possible during the imaging phase. To minimize bias during cell count, concealment of the group allocation was performed (in sham/stroke). None of the animals were excluded after MRI-based inclusion. We did not observe mortality.

## Author Contribution

T.P. designed the study, performed experiments, analyzed results and wrote the manuscript. I.H., S.M., J.L., and M.D. provided technical support. C.H. designed the study, provided resources and reviewed the manuscript. H.G. designed the study, provided resources, acquired funding and reviewed the manuscript.

## Abbreviations

ACA: Anterior Cerebral Artery

dMCAO: distal Middle Cerebral Artery Occlusion

E: Embryonic day

ECs: Endothelial Cells

ICAM: ICAM2

MCA: Middle Cerebral Artery

P: Postnatal day

## Acknowledgments

We thank all members of the Gerhardt lab for discussions and comments, Wolfgang Giese for text comments. We also thank Marie Altmann and the Mouse Facility staff at the MDC for excellent animal care. We gratefully acknowledge Ralf Adams for sharing the BmxCreERT mouse strain and Sagrario Ortega and Taija Makinen for sharing the Vegfr3CreERT mouse strain.

