## Supplementary Material for "Pial collaterals develop through mosaic colonization of capillaries by arterial and microvascular endothelial cells"

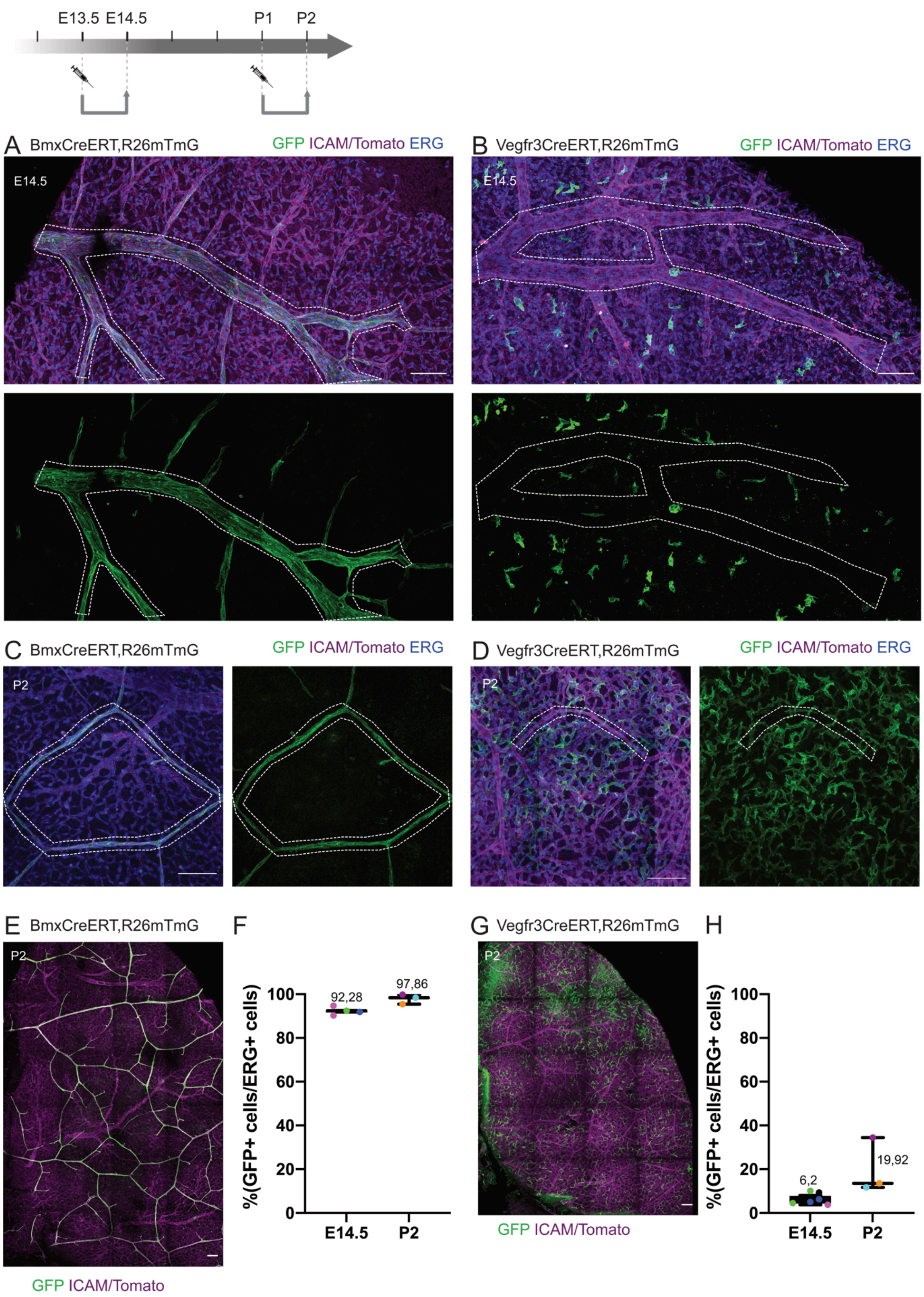


**Figure S1: Bmx-iCreERT and Vegfr3-iCreERT are suitable lineage-tracing tools for pial**

**vasculature**. To test the recombination efficiency, samples were collected 24h after tamoxifen injections (Ez`13.5 - E14.5 or P1 - P2) Scale bar: 100μm. GFP (green), ICAM/Tomato (magenta), ERG (blue). A. E14.5 BmxCreERT;R26mTmG MCA, showing the arterial labeling (GFP channel shown separately below). B. E14.5 Vegfr3CreERT;R26mTmG MCA, showing the venous plexus labeling (GFP channel shown separately below). C. P2 BmxCreERT;R26mTmG, showing the labeling of arterioles and collaterals. D. P2 Vegfr3CreERT;R26mTmG, showing the mosaic labeling of capillaries. E. P2 BmxCreERT;R26mTmG overview (whole hemisphere) image, showing the labeling of arterial vasculature. F. Percentage of the MCA at E14.5 and arterial/collateral circulation at P2 that express Bmx (GFP+ERG+/ERG+ ECs). G. P2 Vegfr3CreERT;R26mTmG overview (whole hemisphere) image, showing the labeling of the plexus ECs. H. Percentage of the plexus ECs at E14.5 and at P2 that express Vegfr3 (GFP+ERG+/ERG+ ECs). Mean and SD are calculated using average data points per animal. Each data point represents a single imaged collateral zone. N≥3 animals, data points from individual animals are color-coded.

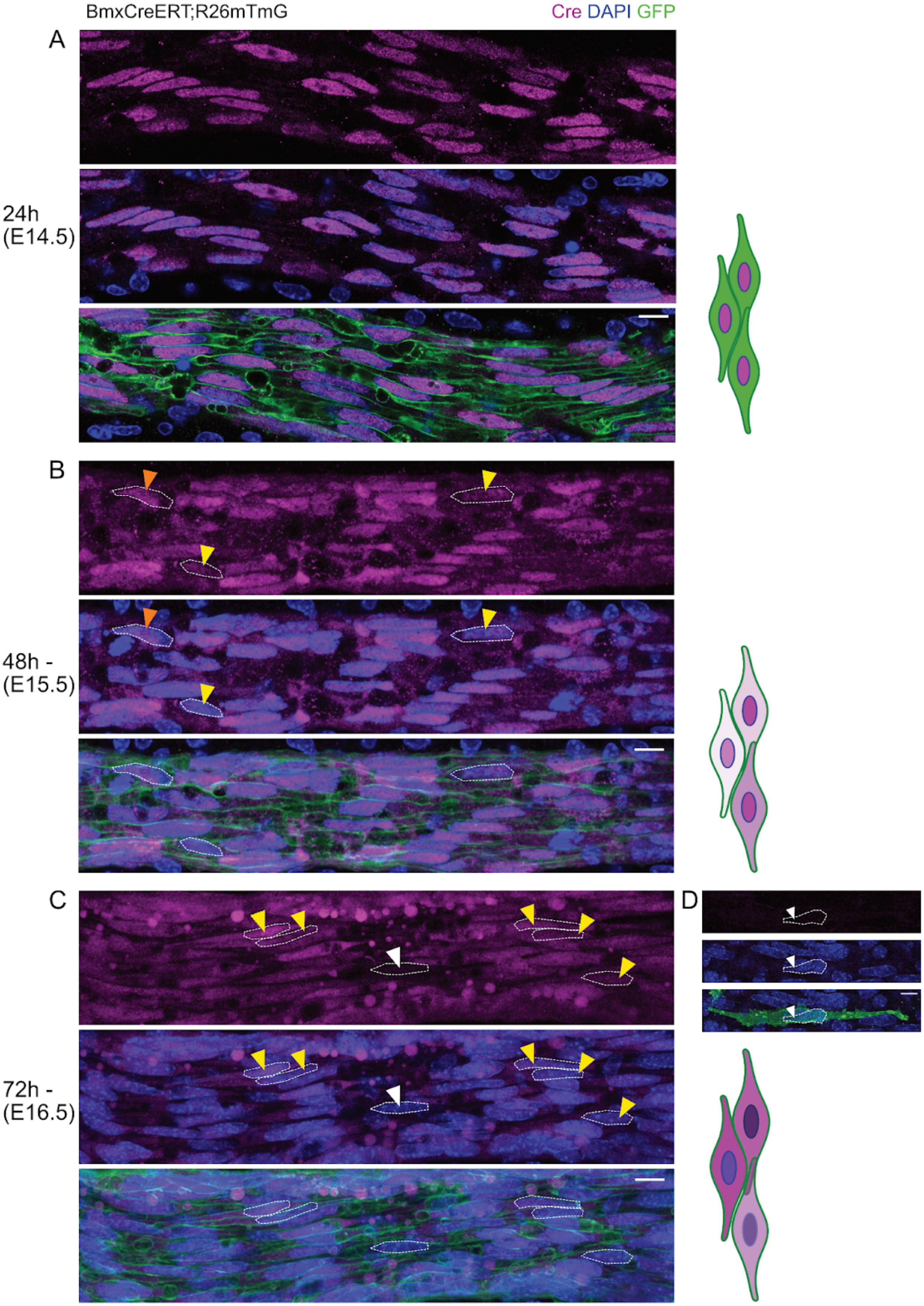
BmxCreERT;R26mTmG
Cre DAPI GFP
A
24h
(E14.5)
B
48h -
(E15.5)
C
72h -
(E16.5)
D

**Figure S2: BmxCreERT recombination window only occurs during 48h upon tamoxifen administration**. Circle of Willis (posterior communicating arteries) of E14.5-E16.5 BmxCreERT;R26mTmG brains, stained against ERα to analyze the distribution of the CreERT protein (under the Bmx promoter) along development (A - C). D. E16.5 BmxCreERT;R26mTmG collateral ECs. White dotted lines represent nuclei of arterial (A-C) and collateral (D) ECs. Orange arrowheads point to GFP-positive/nuclear CreERT; yellow arrowheads point to GFP-positive/nuclear and cytosolic CreERT; white arrowheads point to GFP-positive/CreERT negative cells. Right to A-C: Schematic representation of GFP and CreERT signals detected in panels on the left. At E14.5, ERα localizes inside the nuclei of Bmx-expressing endocardial cells. We didn't observe CreERT expression in forming collaterals, indicating that their GFP labeling is because they are derived from the Bmx-lineage arterial ECs.

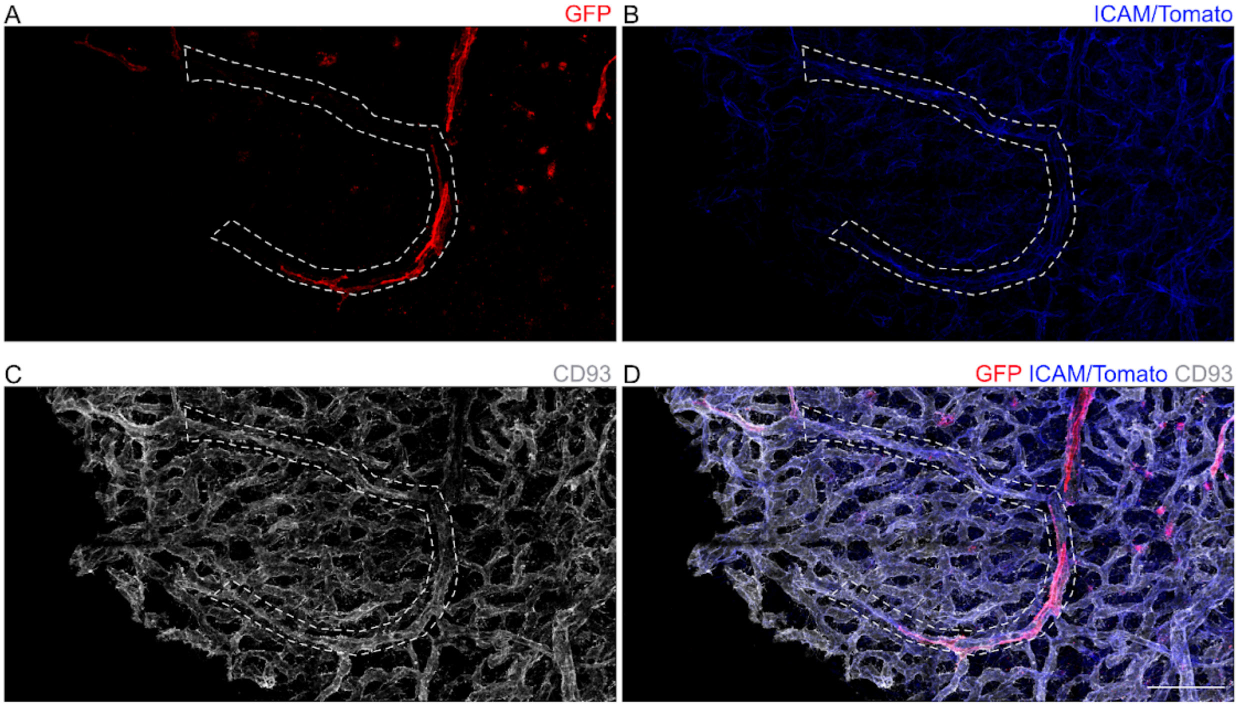


**Figure S3: Pre-collateral vessel structures are characterized by a direct connection to the arterioles, higher diameter than the rest of the plexus, stronger ICAM2-signal and specifically recruit Bmx+ and Vegfr3+ cells.** Bmx-lineage ECs were traced from E13.5 - E16.5. Segmented vessel structure (white dotted lines) is a pre-collateral, shown by the increased ICAM2 positivity (B), direct connection to the upstream arteries and wider diameter than the underlying plexus (C). Bmx-lineage cells (GFP-red) are colonizing the pre-collateral (A,D).

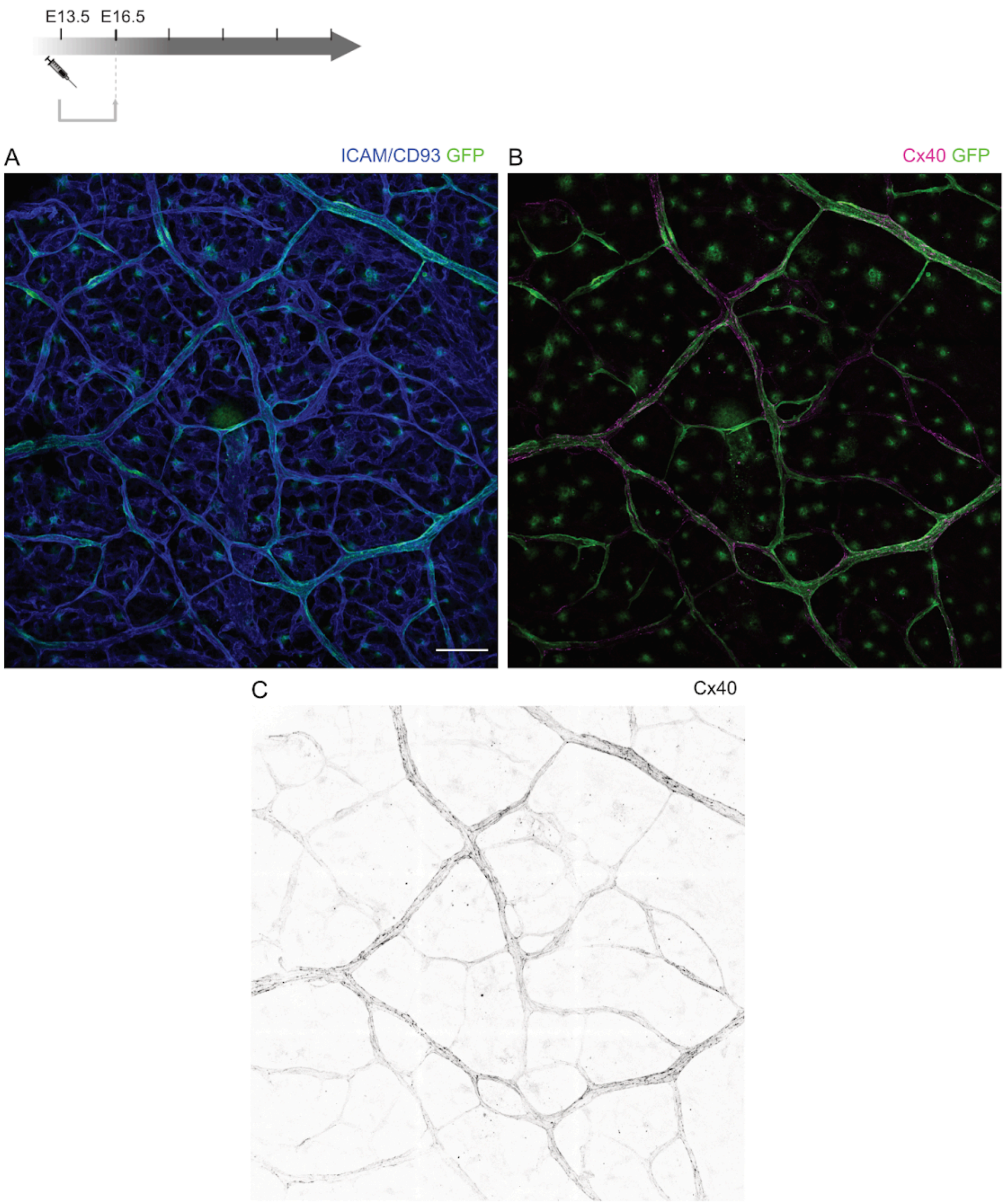


**Figure S4: Connexin40 (Cx40) immunostaining shows a zonation pattern: strongest in biggest arteries, weakest/absent in nascent collaterals**. Bmx-lineage ECs were traced from E13.5 - E16.5 (top left scheme). 16.5 BmxCreERT;R26mTmG pial pre-collateral zone (including bigger arteries/arterioles). A. ICAM2/CD93 - blue, GFP - green. B. Cx40 - magenta, GFP - green. C. Cx40 - gray.

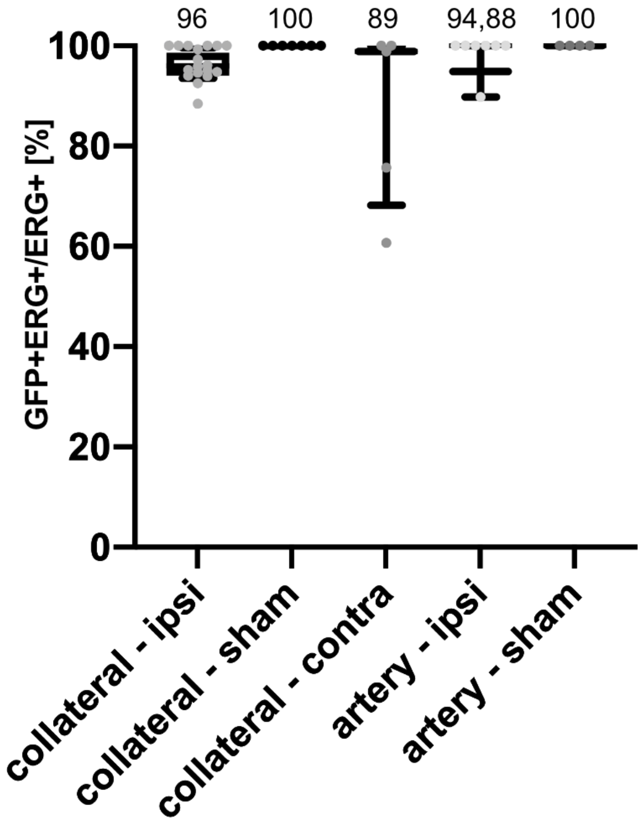


**Figure S5: Percentage of Bmx-derived GFP+ ECs in collaterals after dMCAO/sham operation.** Bmx-Cre mTmG animals were injected at P1 and underwent either stroke or sham operation in adulthood. N≥3 animals, each dot representing an average of one collateral vessel. Mean ±SD. ipsi - ipsilateral, contra - contralateral. N=2 animals for sham (technical control) and N=5 for stroke animals. Mean and SD are calculated using average data points per animal.

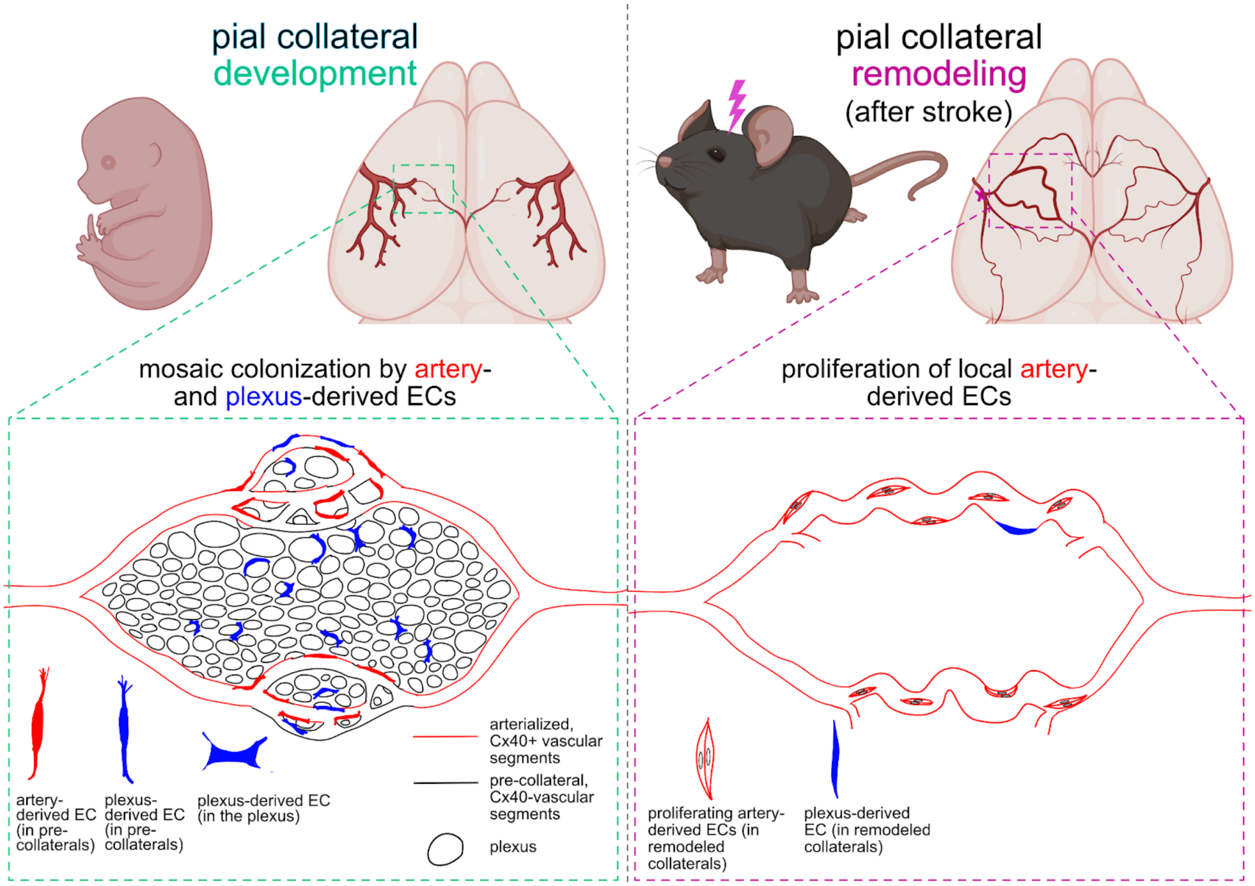


**Figure S6: Graphical Abstract.** A visual summary of our main findings. On the left side, the updated model of embryonic pial collateral development: in red lines are arterialized, Cx40+ vascular segments (already established arteries and collaterals), black lines indicate pre-collateral, Cx40- vascular segments, which are being populated by ECs of arterial (red ECs) and microvascular (blue, elongated ECs) lineage in the process we coined *mosaic colonization*. There is a dense, underlying microvascular plexus intercalated with the pre-collateral zone. The right side of the abstract summarizes our model of collateral remodeling after stroke: we found that this process in part depends on proliferation of local ECs of arterial lineage (elongated, red-outlined sister ECs with white nuclei), and does not involve proliferation of local (blue elongated ECs) or newly recruited microvascular ECs (not shown).
